# Competition and Cooperation of Assembly Sequences in Recurrent Neural Networks

**DOI:** 10.1101/2023.11.03.565506

**Authors:** Tristan M. Stöber, Andrew B. Lehr, Marianne Fyhn, Arvind Kumar

## Abstract

Neural activity sequences are ubiquitous in the brain and play pivotal roles in functions such as long-term memory formation and motor control. While conditions for storing and reactivating individual sequences have been thoroughly characterized, it remains unclear how multiple sequences may interact when activated simultaneously in recurrent neural networks. This question is especially relevant for weak sequences, comprised of fewer neurons, competing against strong sequences. Using a non-linear rate model with discrete, pre-configured assemblies, we demonstrate that weak sequences can compensate for their competitive disadvantage either by increasing excitatory connections between subsequent assemblies or by cooperating with other co-active sequences. Further, our model suggests that such cooperation can negatively affect sequence speed unless subsequently active assemblies are paired. Our analysis, validated by an analytically tractable linear approximation, characterizes the conditions for successful sequence progression in isolated, competing, and cooperating sequences, and identifies the distinct contributions of recurrent and feed-forward projections. This proof-of-principle study shows how even disadvantaged sequences can be prioritized for reactivation, a process which has recently been implicated in hippocampal memory processing.

## 2 Introduction

Sequences of neural activity are a universal phenomenon in the brain, fundamentally underpinning a range of functions including olfactory processing (Friedrich and Laurent, 2001), birdsong generation (Hahnloser et al., 2002), motor control (Eichenlaub et al., 2020), and episodic memory encoding in the hippocampus (O’Keefe, 1976; Dragoi and Buzsáki, 2006; Foster and Wilson, 2006; Diba and Buzsáki, 2007). These sequences unfold over various timescales and can be driven by either external stimuli or intrinsic mechanisms. Thus, to understand information processing in the brain, we need to comprehend the dynamics of neural activity sequences.

The emergence and reliable propagation of individual neural activity sequences have been extensively studied using computational models (Amari, 1977; Arnoldi and Brauer, 1996; Hertz, 1997; Diesmann et al., 1999; Abeles et al., 2004; Kumar et al., 2008; York and van Rossum, 2009; Fiete et al., 2010; Itskov et al., 2011; Lu et al., 2011; Azizi et al., 2013; Kappel et al., 2014; Chenkov et al., 2017; Murray et al., 2017; Seeholzer et al., 2019; Spreizer et al., 2019; Michaelis et al., 2020; Maes et al., 2020b,a; Spalla et al., 2021; Lehr et al., 2023). A number of studies characterized conditions for storing and reactivating multiple sequences in recurrent networks(Arnoldi and Brauer, 1996; Abeles et al., 2004; Kumar et al., 2008; Azizi et al., 2013; Maes et al., 2020a; Spalla et al., 2021; Lehr et al., 2023). However, interactions between sequences within a network are less understood, and in particular the influence of competition and cooperation on sequence reactivation has not yet come into focus.

A popular experimental paradigm to expose the functional role of neural activity sequences is to record the activity of hippocampal neurons in a spatial navigation task, commonly performed in rats or mice. While traversing an environment, place cells are activated in a sequential manner (O’Keefe, 1976; Dragoi and Buzsáki, 2006). Subsequently, when the animal is resting, planning or consuming, the same neural activity sequences may be reactivated (or replayed) at a faster time scale during sharp wave ripple (SPWR) events (Wilson et al., 1994; Skaggs and McNaughton, 1996; Ji and Wilson, 2007; Diba and Buzsáki, 2007). Such offline reactivation can represent multiple distinct experiences (Silva et al., 2015). However, it is assumed that normally only one sequence is reactivated per sharp-wave ripple (He et al., 2020).

Replay of sequences is crucial for memory consolidation (Girardeau et al., 2009; Dupret et al., 2010; Fernández-Ruiz et al., 2019; Oliva et al., 2020). Successful generation of long sequences during SWR is associated with better memory (Fernández-Ruiz et al., 2019). Moreover, the probability that a particular sequence will be reactivated varies with experience, with novel and reward-related sequences being prioritized (McNamara et al., 2014; Igata et al., 2021; Singer and Frank, 2009; Ambrose et al., 2016, but see Gupta et al., 2010). Intriguingly, the fact that neither the generation of sharp-wave ripples (Bragin et al., 1995; Yamamoto and Tonegawa, 2017) nor the reactivation of sequences (Chenani et al., 2019) are abolished by lesions or inhibition of the medial entorhinal cortex, the primary input structure to the hippocampus, suggests the existence of inherent mechanisms for sequence prioritization within the hippocampus.

Hippocampal activity sequences differ in key properties depending on which information they represent. When encoding the location of objects and other animals fewer hippocampal place cells are recruited and their firing rates are lower compared to place cells for the animal’s own location (Danjo et al., 2018; Omer et al., 2018). Thus, hippocampal sequences are likely composed of differently sized cell assemblies. In the following we call sequences with large assemblies strong and those with small assemblies weak. To consolidate their corresponding experiences, it is conceivable that both weak and strong sequences compete for reactivation during SPWRs.

A computational model suggests that successful reactivation becomes more difficult for weak sequences, unless recurrent connections within and/or feed-forward projections between cell assemblies are strengthened (Chenkov et al., 2017). However, the required amount of potentiation increases non-linearly with decreasing assembly size, and synapses may quickly reach their physiological boundaries (Chenkov et al., 2017). If multiple sequences are activated at the same time, mutual inhibition between them may create a winner-take-all type competition. In such a scenario, weak sequences essentially stand no chance of winning the competition.

Here, we explore how weak sequences may cooperate to win over stronger sequences during replay events. Inspired by recent findings about gated synaptic plasticity and mutual feed-forward inhibition between region CA3 and CA2 in the hippocampus, we proposed that co-occurring sequences in these regions may be selectively paired by the release of neuromodulatory substances (Stöber et al., 2020). In addition to linking distinct information (Mankin et al., 2015; Lee et al., 2015; Wintzer et al., 2014) in each region, mutual excitatory support between CA3 and CA2 sequences may ensure their reactivation, while at the same time recruiting sufficient inhibition to suppress competing sequences (He et al., 2020; Lehr et al., 2021).

To develop a theoretical understanding based on these hippocampal insights, we demonstrate that cooperation and competition of assembly sequences can be implemented in a rate-based model. Within a sequence, reliable and fast signal transmission is achieved by excitatory feed-forward projections between subsequent assemblies, employing balanced amplification (Murphy and Miller, 2009; Chenkov et al., 2017). Competition and cooperation are implemented by feed-forward inhibition and excitation across assemblies. Characterizing conditions for competition and cooperation, we show that a) feed-forward excitation is crucial, but must remain within a certain range to avoid excessive and persistent activation, b) recurrence within assemblies helps the surviving sequence to recover, c) feed-forward inhibition can mediate competition, d) excitatory coupling between co-active assemblies allows weak sequences to win, but slows sequence progression, and e) preferentially pairing subsequently instead of co-active assemblies maintains sequence speed. Taken together, these results demonstrate that reactivation dynamics of neural sequences are shaped both by modifying feed-forward properties as well as by interactions among multiple sequences.

## 3 Results

### 3.1 Conditions for progression of a single sequence

We used a rate-based model with a non-linear activation function to first study the progression of a single assembly sequence (Fig. 1a,b). Each assembly is composed of discrete and recurrently interacting populations of excitatory and inhibitory neurons. Sequences are defined by connecting subsequent excitatory populations with feed-forward projections. In addition, all assemblies – independent of their position in the sequence – send feed-forward inhibition to each other; they send excitatory projections to each other’s inhibitory populations. To start the sequence the excitatory population of the first assembly receives external stimulation. To characterize successful sequence progression, we defined four conditions: 1) All active: Within each assembly, the excitatory population must be activated at least at one point in time. 2) All informative: In addition, each excitatory population must exceed the activity of others at least one point in time. 3) Sparse activity: Global activity of the whole network must be sparse, e.g. peak activity is not to be reached by more than two assemblies at any point in time. 4) Order: Peak activation of any excitatory population must maintain its predefined order.

**Figure 1:**
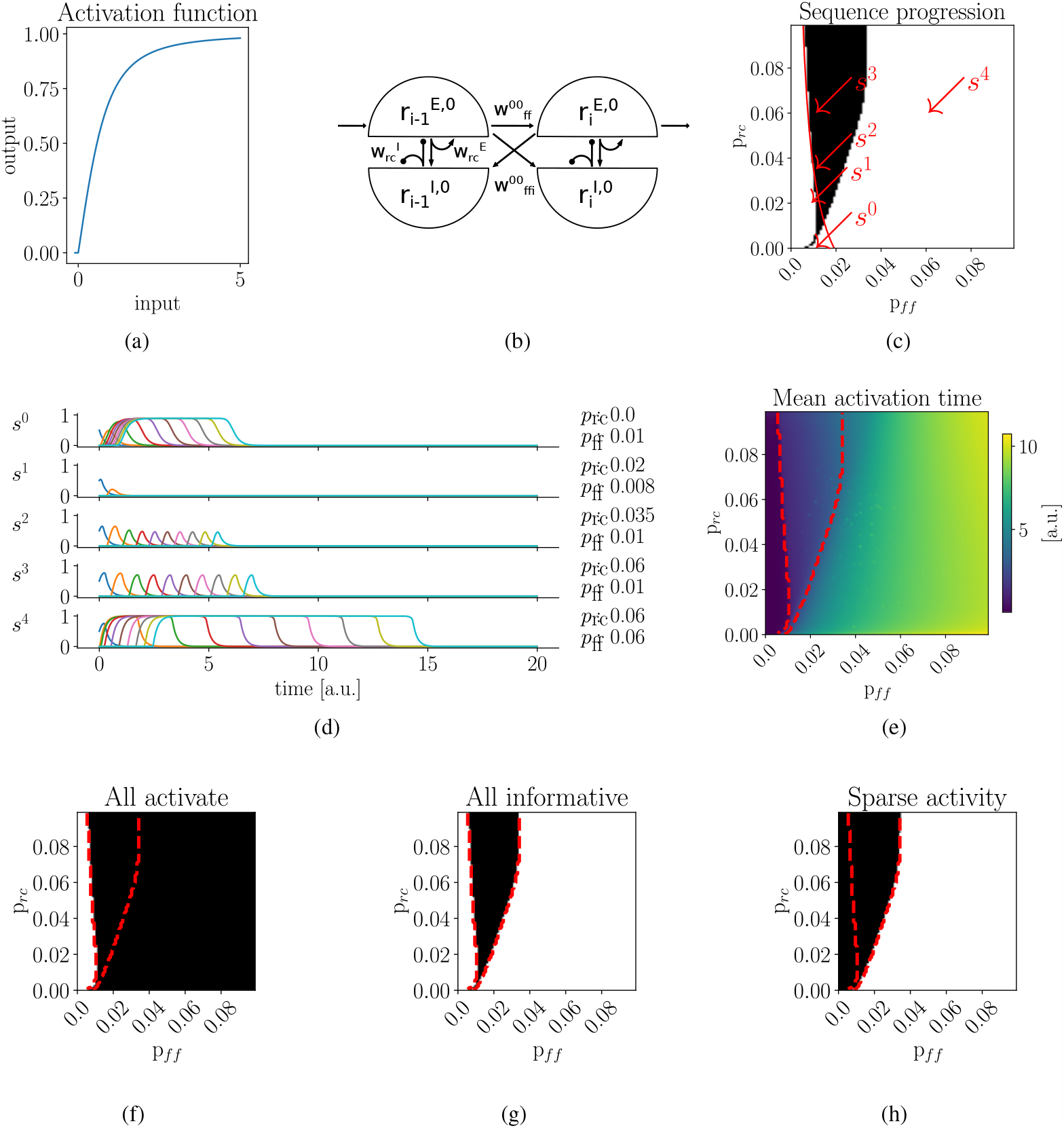
Recurrent and feed-forward interactions influence the progression of a single sequence. **a)** Nonlinear activation function of excitatory and inhibitory populations. **b)** Connections within and between assemblies in a single sequence. Each assembly is formed by two recurrently interacting populations, representing excitatory (E) and inhibitory (I) neurons. A given sequence *s*^0^ is established by connecting subsequent excitatory populations via feed-forward excitatory projections with strength 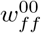 . Assemblies generally suppress each other via feed-forward inhibition; excitatory projections from excitatory to all inhibitory populations of other assemblies with strength 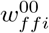 . **c)** Successful sequence progression, black region, depends on both recurrent, *p*_*rc*_ and, feedforward, *p*_*ff*_, connection probability. Red line, analytic solution of the linearized rate model for sustained activity propagation. For low values of *p*_*rc*_, the non-linear rate model and the analytic solution diverge. **d)** Example sequences, corresponding to red arrows in b). Only *s*^2^ and *s*^3^ successfully reactivate. Activity 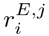 of every third excitatory population is shown. Different colors correspond to different excitatory populations. **e)** Mean activation time across all assemblies. Large values of *p*_*ff*_ lead to persistent activity and, thus, to large mean activation time. **f)** Parameter region (black) where all excitatory populations in sequence become activated, fulfilling condition 1: *All active*. **g)** Parameter region fulfilling condition 2: *All informative*, all excitatory populations must exceed activity of others at least once (black). **h)** Parameter region fulfilling condition 3: *Sparse activity*. Red, dashed contours in c), e), f) and g) correspond to black region for successful sequence progression in b).

Successful sequence progression depends on both recurrent and feed-forward projections. The strength of each connection is a product of the respective excitatory or inhibitory population size, *M* ^*{E,I}*^, synaptic strength, *g*^*{E,I}*^ and recurrent or feed-forward connection probability, *p*_*{rc,ff}*_. To investigate the dependence of sequence progression on connection strength, we systematically varied *p*_*rc*_ and *p*_*ff*_, simultaneously for excitatory and inhibitory projections. We found that the parameter region allowing successful sequence progression for *p*_*ff*_ is relatively narrow compared to *p*_*rc*_ (Fig. 1c, black region). Closer investigation revealed that, without sufficient feed-forward projections, activity dies out (*s*^1^ in Fig. 1d), preventing all assemblies from being activated (Fig. 1f), violating condition 1 (Fig. 1g). By contrast, strong feed-forward projections led to rapid and persistent activation (*s*^4^ in Fig. 1d, 1e), violating the condition 2 (Fig. 1h). However, if excitatory and inhibitory populations recurrently interact with sufficient strength, assembly activation can become transient, allowing sequences to progress in a sparse fashion for an increasing range of feed-forward weights, condition 3 (*s*^2^ and *s*^3^ in Fig. 1d).

The V-shape of the parameter region reflecting successful progression illustrates the dual role of recurrent interactions. On its left flank (for weak feed-forward connections), increasing recurrent interactions, *p*_*rc*_ *>* 0.025, decreases the required feed-forward weights, *p*_*ff*_ by positively amplifying weak inputs. On the right flank (for strong feed-forward connections), stronger recurrent inhibition prevents persistent activity and, thus, increases permissible feed-forward weights.

Progression of single assembly sequences can be approximated by an even simpler linear dynamical system (Chenkov et al., 2017). Under the assumption of stationarity, we analytically determined the minimal value of *p*_*rc*_ required for sustained activity in subsequent assemblies in relation to *p*_*ff*_ (for details see Methods). We show the simulation results are in close agreement to the analytically determined values of *p*_*rc*_ and *p*_*ff*_ for *p*_*rc*_ *>* 0.025 (Fig. 1c, red line). For *p*_*rc*_ *<* 0.025 the analytical approximation diverges from the simulation results (see Discussion). For very low recurrence values, *p*_*rc*_ ∼ 0, sequence progression is limited to few specific values of *p*_*ff*_ (*s*^0^ in Fig. 1d, 1e). Note that the solutions of the the linear approximation are influenced by a scaling factor *c*, related to the slope of the neuron’s input-output function. Throughout the article, we retain *c* = 0.163, as determined by solving for *c* under the parameters of example sequence *s*^2^ (see Methods).

### 3.2 Competition between two sequences

Next, we studied competition between two sequences, *s*^0^ and *s*^1^. As before, each assembly sends feed-forward inhibition to all other assemblies, both within and between sequences (Fig. 2a). If the first assemblies in both sequences are simultaneously activated, the interplay between excitation within the assemblies and inhibition between sequences can lead to one of four scenarios: a) Activity in both sequences ceases before the sequence is completed; referred to as *no winner*; b) *s*^0^ successfully progresses and *s*^1^ ceases; referred to as *s*^0^ *wins*; c) *s*^1^ successfully progresses and *s*^0^ ceases; referred to as *s*^1^ *wins*; d) both sequences successfully progress; referred to as *both win*.

**Figure 2:**
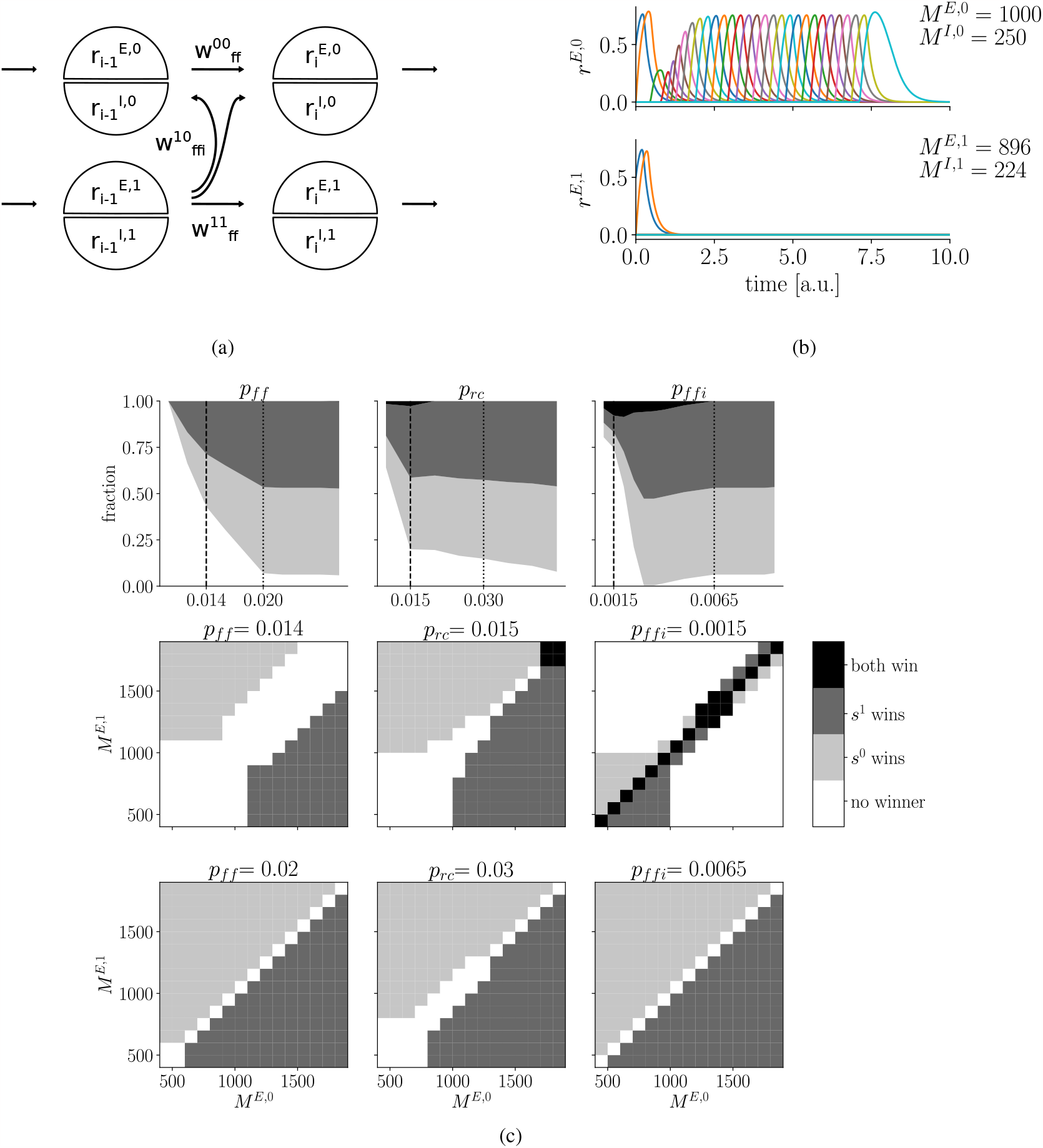
Competition between two sequences. **a)** Scheme of connections for competition scenario. Two sequences, *s*^0^ and *s*^1^, compete via feed-forward inhibition between all assemblies. For visual clarity, only feedforward inhibition with strength 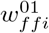 from one assembly of *s*^0^ to two assemblies of *s*^1^ is shown. **b)** Example: Larger sequence *s*^0^ wins over *s*^1^. Only activities of excitatory populations are shown. Both sequences suppress each other’s activity until *s*^1^ ceases and *s*^0^ recovers. Colors repeat after 10 assemblies. **c)** Competition scenarios for different values of feed-forward, recurrent, and feed-forward inhibition connection probability. *p*_*ff*_, *p*_*rc*_, *p*_*ffi*_ are separately varied. Middle and bottom rows exemplify competition outcomes when scanning the *M*^*E*,0^, *M*^*E*,1^ space for different values of *p*_*ff*_, left, *p*_*rc*_, center, or *p*_*ffi*_, right. Upper row summarizes the distribution of competition scenarios for each scan of the *M*^*E*,0^, *M*^*E*,1^ space. Specific values of low and high value examples are indicated by dashed and pointed lines, corresponding to upper and center rows.

To exemplify competition dynamics, we let two sequences compete for a given set of parameters (Fig. 2b). After an initial surge, activity diminished in both sequences. While *s*^1^ ceased, activity in *s*^0^, with slightly larger assemblies, recovered and successfully progressed.

The competition outcome depends on assembly sizes as well as interactions within and between sequences. To systematically characterize the occurrence of the four competition scenarios, we varied assembly sizes *M*^*E*,0^, *M*^*E*,1^ for different connection probabilities of either feed-forward excitation *p*_*ff*_, recurrent excitation *p*_*rc*_, or feed-forward inhibition *p*_*ffi*_. Note, in the following, sizes of the inhibitory populations, *M*^*I*,0^, *M*^*I*,1^, are scaled accordingly to maintain a constant ratio of excitatory and inhibitory population sizes.

For example, for moderate levels of feed-forward excitation, *p*_*ff*_ = 0.014, relatively large assemblies, *M*^*E*,0^, *M*^*E*,1^ *>* 1400 were required for one sequence to win over the other (Fig. 2c, central row in left column). Nevertheless, even for large assemblies, the difference between sequences had to be prominent, otherwise both sequences ceased to exist. By contrast, if feed-forward excitation is increased, even moderately sized assembly sequences could win as long as they are larger than their competitor (Fig. 2c, bottom row in left column). As a consequence of this, the fraction of the *M*^*E*,0^, *M*^*E*,1^ parameter space spanned by either *s*^0^ or *s*^1^ winning increased with a rise in the strength of feed-forward connections, *p*_*ff*_, until it hit an upper bound (Fig. 2c, upper row in left column).

Without recurrence, even sequences with large assemblies failed to successfully propagate when competing. As we showed in Figure 1c and know from the literature on synfire chains (Hertz, 1997; Diesmann et al., 1999; Kumar et al., 2010), individual sequences can progress without recurrent interactions. However, we hypothesized that in a competition scenario, recurrence is paramount for the surviving sequence to recover. To test this, we characterized the competition outcome for a range of assembly sizes given different values of *p*_*rc*_. Consistent with our expectation, for relatively weak recurrence, *p*_*rc*_ = 0.015, larger assemblies were required to avoid that both sequences cease their progression (Fig. 2c, central row in central column). Surprisingly, we found that weak recurrence allows both strong sequences to win (Fig. 2c, black region, central row in central column). With an increase in the recurrence, *p*_*rc*_ = 0.03, the fraction of the *M*^*E*,0^, *M*^*E*,1^ parameter space spanned by either *s*^0^ or *s*^1^ winning increased (Fig. 2c, bottom row in central column). Thus, we conclude that recurrence is indeed crucial when sequences compete.

Feed-forward inhibition ensures that only one sequence wins. Here, sequences competed by inhibiting each other. Therefore, we expected that relatively weak feed-forward inhibition will allow both sequences to win. Again classifying competitions outcomes, we could indeed show that for low values of *p*_*ffi*_ a considerable fraction of the *M*^*E*,0^, *M*^*E*,1^ parameter space was covered by the *both win* scenario (Fig. 2c, black region, upper and central row, right column). Further, we found that weak feed-forward inhibition corresponded to a large fraction of failed progressions for both sequences, *no winner*. Without inhibition between assemblies of the same sequence, activity in each cell assembly became persistent, violating the sparsity condition (data not shown). On the other hand, when *p*_*ffi*_ was increased, the *both win* case disappeared (Fig. 2c, upper and bottom row, left column).

Compensating for small assemblies by increasing feed-forward weights may become physiologically implausible. As proposed in (Stöber et al., 2020), a weak sequence, comprised of smaller assemblies, competing with a strong sequence, can ensure progression by further potentiating feed forward weights. However, we hypothesized that the required amount of potentiation scales non-linearly with assembly size – as already shown for individual sequences (Chenkov et al., 2017) – and therefore, may hit physiological boundaries for weak sequences. We explicitly tested this prediction by varying the respective parameters for sequence *s*^1^; keeping parameters in *s*^0^ fixed. In agreement with the hypothesis, we found a non-linear increase in the required feed-forward connection probability 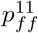 for decreasing assembly sizes *M*^*E*,1^ (Fig. 3a). If both assembly sizes and feed-forward weights were strong (*no winner region* in upper right corner of Fig. 3a), persisting activity violated both the activation and the sparsity condition (data not shown).

**Figure 3:**
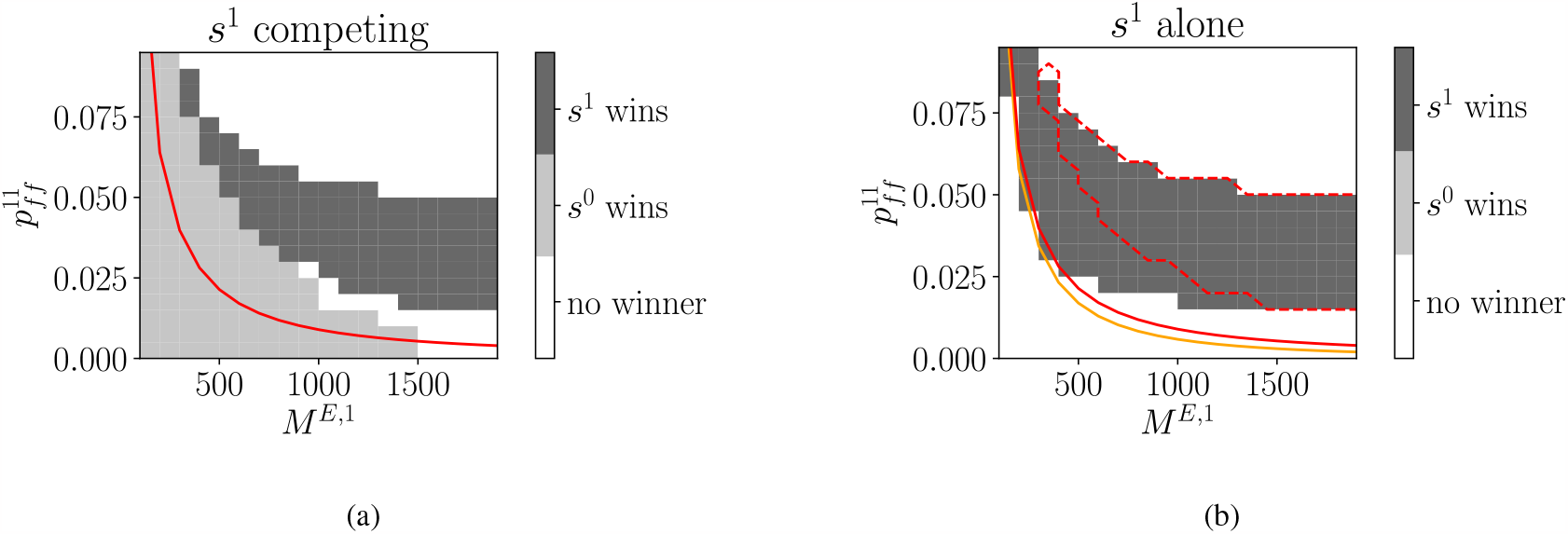
Competition makes it harder for sequences with small assemblies to ensure progression by strengthening feed-forward weights. **a)** For fixed parameters of *s*^0^, *M*^*E*,0^ = 1000, 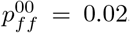, assembly size *M*^*E*,1^ and feed-forward connection probability 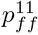 of *s*^1^ are varied. For *s*^1^ to win (dark area), smaller assemblies must be compensated by increasingly larger feed-forward weights. Red line, analytic prediction of linearized rate model for sustained activity propagation in *s*^1^ with constant activity in *s*^0^, 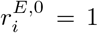. **b)** Same as a), but without activating *s*^0^. Orange line, analytic prediction for sustained activity propagation in *s*^1^ with *s*^0^ silenced. Red dashed contour line reflects *s*^1^ *wins* area and red line analytic prediction from competition scenario in a).

To gain an analytic understanding of the competition scenario, we extended the simplified linear rate model to include a second sequence. Keeping the activation of excitatory populations in *s*^0^, we solve for 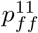, the minimal required feed-forward weight to ensure sustained activity of *s*^1^ (Fig. 3a, red line, see Methods). However, the analytic solution predicts lower required weights compared to the rate-model simulation. One potential explanation for this difference may be that feed-forward inhibition in the rate model is between all assemblies, while it is only to the next competing assembly in the simplified linear model (see Discussion).

To compare the presented results to a situation without competition, we repeated the simulation and analytic calculations with silenced *s*^0^ (Fig. 3b). As before, the required feed-forward connection probability increased non-linearly with decreasing assembly size. However, without a competing sequence, also smaller feed-forward connection probabilities allowed successful propagation. This holds true for both the analytical prediction and the simulation results. In conclusion, these findings show that competition increases the required strength of feed-forward weights, making it even more difficult to reactivate sequences with small assemblies.

### 3.3 Cooperation and competition between three sequences

Given the physiological limits on the potentiation of feed-forward projections, an alternative or additional way for sequences to ensure progression despite competition is to mutually support each other. This may happen if simultaneously active assemblies in co-occurring sequences are paired by Hebbian plasticity (Stöber et al., 2020). To demonstrate both cooperation and competition between assembly sequences, we created a minimal scenario with one strong, and two weak sequences (Fig. 4a). As before, all assemblies mutually inhibited each other and the excitatory populations at the start of each sequence were simultaneously activated. The strong sequence *s*^0^ has a competitive advantage due to its larger assemblies. As expected, without any cooperation between *s*^1^ and *s*^2^, sequence *s*^0^ won (case *c*_0_, Fig. 4b, Fig. 4c).

**Figure 4:**
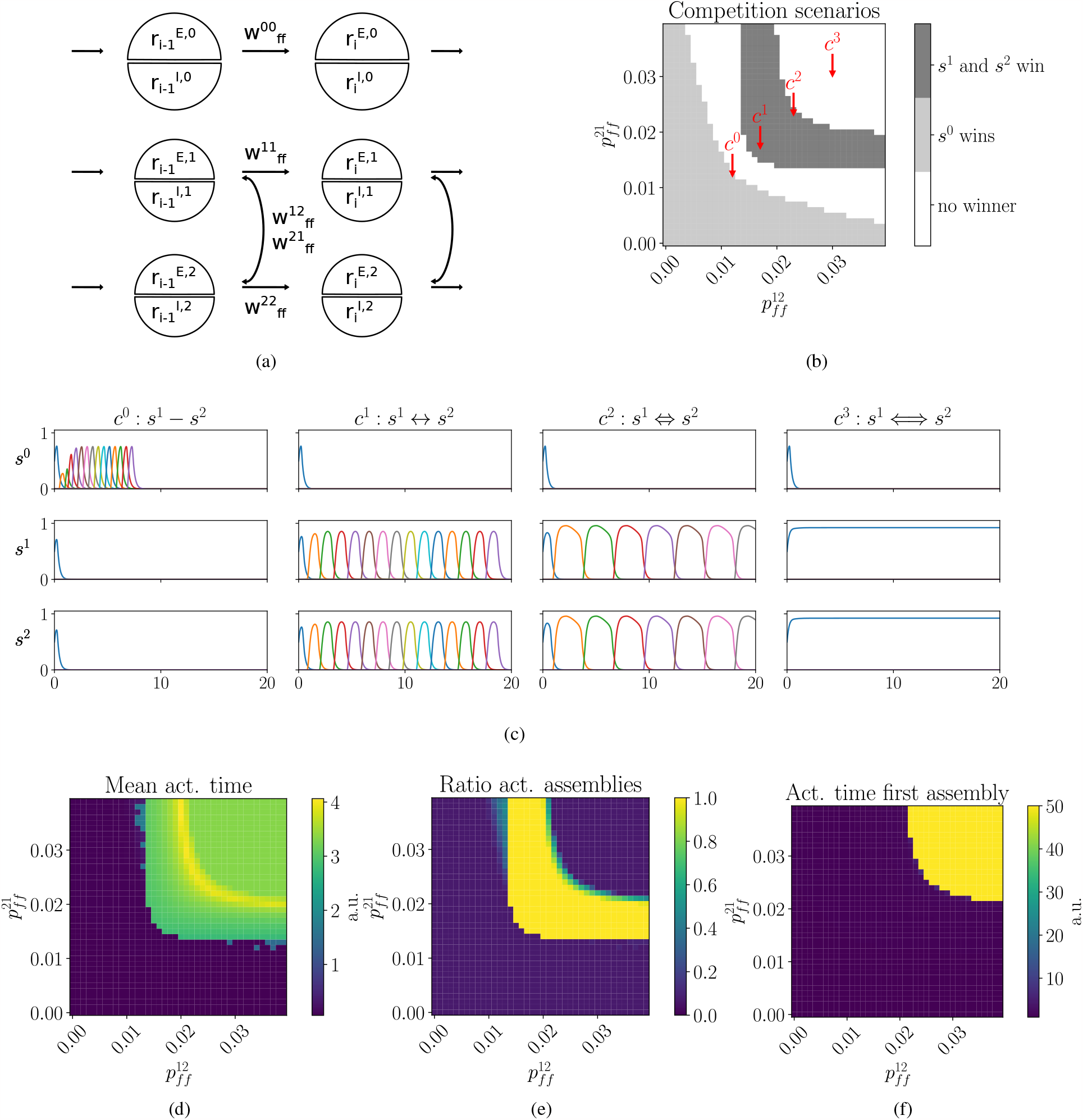
Cooperation via mutual excitation between assembly sequences. **a)** Network scheme for competition and cooperation between discrete assembly sequences. Sequence with larger assemblies, *s*^0^, competes with *s*^1^ and *s*^2^. Competition via feed-forward inhibition between all sequences not shown. Cooperation between *s*^1^ and *s*^2^ through reciprocal excitatory connections to co-active assemblies with strength 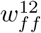 and 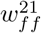 . Larger assemblies of *s*^0^ are indicated by larger circles. **b)** Connection probabilities between *s*^1^ and *s*^2^, 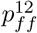 and 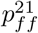, are varied. Sufficiently strong mutual excitation is required for *s*^1^ and *s*^2^ to outcompete *s*^0^. **c)** Examples: *c*_0_ : *s*^1^ − *s*^2^, mutual excitatory interactions between *s*^1^ and *s*^2^ not sufficient; *c*_1_ : *s*^1^ ↔ *s*^2^, pairing between *s*^1^ and *s*^2^ strong enough to win; *c*_2_ : *s*^1^ ⇔ *s*^2^, increased excitatory interactions lead to slower sequence progression, e.g. longer activation times; *c*_3_ : *s*^1^ ⇔ *s*^2^, if excitatory interactions are too strong, sequence progression fails because first assemblies of *s*^1^ and *s*^2^ remain active. Only activity of excitatory populations is shown. **d)** Mean activation times of excitatory populations in *s*^1^. Strong mutual excitation leads to longer activation times and slow sequence progression. **e)** Ratio of activated excitatory populations in *s*^1^. Only in region with successful cooperation with *s*^2^ all assemblies of *s*^1^ are activated. **f)** Activation time of first excitatory population in *s*^1^. Strong interactions halt propagation, because early assemblies maintain activity over the full simulation duration.

When weak sequences were able to cooperate, they could however overcome a strong competitor. We introduced feed-forward excitatory projections between co-active excitatory populations in *s*^1^ and *s*^2^, summarized by their strengths 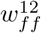 and 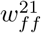 . Given sufficient mutual support, *s*^1^ and *s*^2^ were able to out-compete *s*^0^ (*c*_1_, Fig. 4b, Fig. 4c). However, the stronger the mutual excitatory connections, the longer were the activation times of excitatory populations (*c*_1_ vs. *c*_2_, Fig. 4d). Increasing the excitatory interactions further led to persistent activity in the first assemblies, halting successful sequence progression (*c*_3_, Fig. 4c, 4e, 4f).

### 3.4 Potentiation of excitatory synapses to subsequently active assemblies in paired sequence increases propagation speed

In the previous section we observed that pairing sequences by potentiating co-active assemblies can indeed facilitate their reactivation, but it slows sequence progression and, if too strong, leads to persistent activity. Thus, we hypothesized that sequence speed can be increased by introducing excitatory projections to subsequently active assemblies (see Fig. 5a). Adding this type of projection to the three sequence model and explicitly measuring sequence speed by the inverse of the median interpeak interval of excitatory populations, we observed a range of different speeds depending on the relative levels of potentation between co-active and subsequent assemblies (Fig. 5b,5c). As expected, stronger potentiation between co-active assemblies led to slower progression (purple region, Fig. 5b). Additionally increasing the synaptic strength between subsequent while maintaining strong synapses between co-active assemblies marginally increased speed at the expense of prolonged activation times of individual assemblies (*c*^0^ vs. *d*^0^ and *d*^1^). However, reducing synapse strength between co-active while maintaining relatively strong synapses to subsequent assemblies can increase sequence speed up to the level of the competing sequence *s*^0^ (yellow region, Fig. 5b; example *d*^2^, Fig. 5c). Thus, potentiating subsequently active assemblies can indeed facilitate reactivation of paired sequences while preserving the timescale of sequence progression.

**Figure 5:**
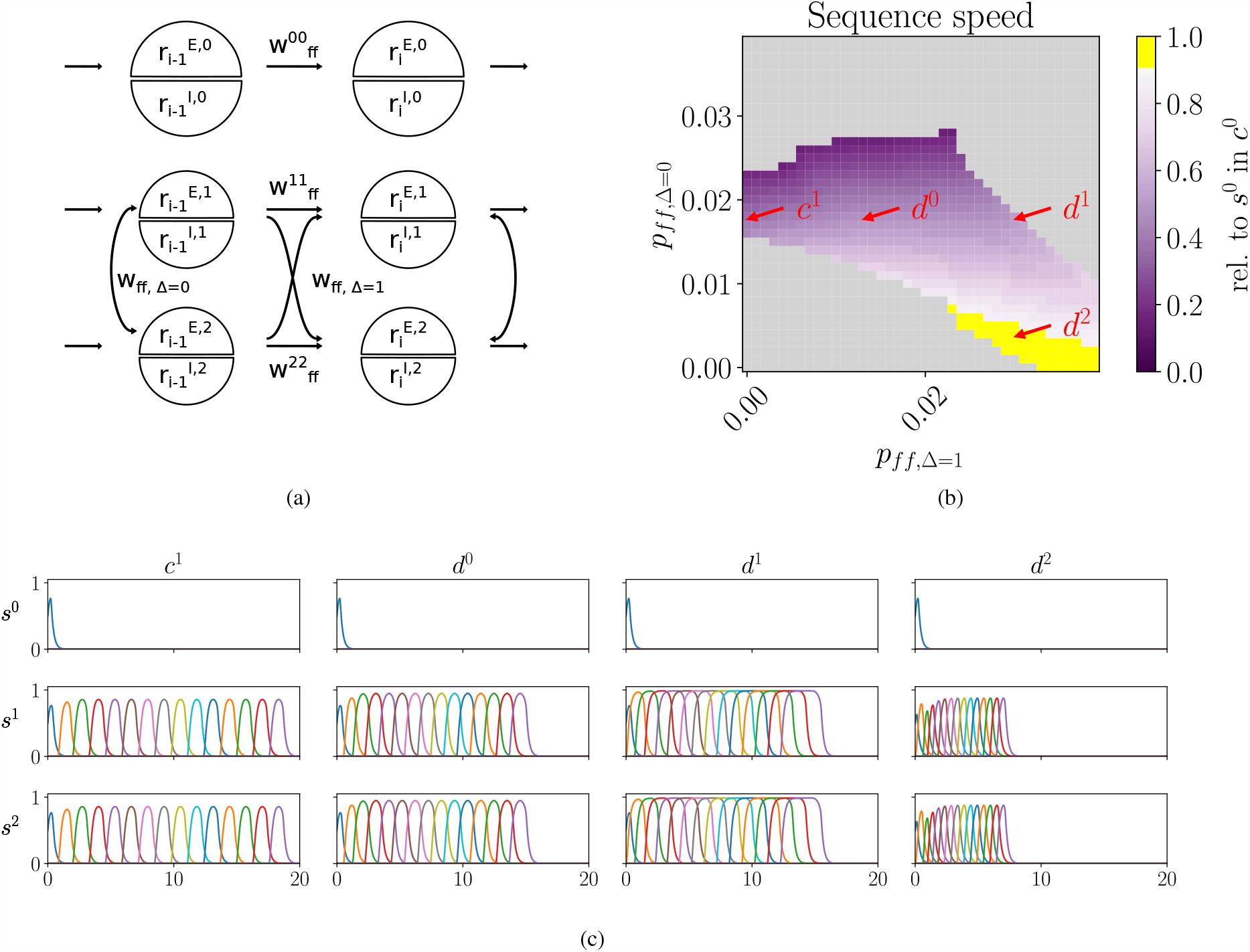
Shifting feedforward excitation from co-active to subsequent assemblies increases speed of cooperating sequences. **a)** Beyond prior simulations which modified feedforward excitation solely between simultaneously active assemblies, *w*_*ff*,∆=0_, we also introduce feedforward excitation to subsequent assemblies, *w*_*ff*,∆=1_, in cooperating sequences *s*^1^ and *s*^2^. **b)** Sequence speed of *s*^1^, relative to *s*^0^ of scenario *c*^0^ from Fig. 4b. Successful reactivations of *s*^1^ and *s*^2^ are shown for reduced (shades of purple) and similar speed as competing sequence (yellow interval, from 0.9-1.). Non-successful reactivations are indicated in grey. Adjustments in weights *w*_*ff*,∆=0_ and *w*_*ff*,∆=1_ are achieved by altering the corresponding connection probabilities *p*_*ff*,∆=0_ and *p*_*ff*,∆=1_. **c)** Examples: *c*^1^ From Fig. 4b without pairing between subsequently active assemblies *d*^0^ Increasing feedforward excitation to subsequent assemblies can counterbalance the reduced speed brought on by the additional feedforward excitation between co-active assemblies. *d*^1^ Further increasing feedforward excitation to subsequent assemblies expands activation duration of individual assemblies. *d*^2^ To reach speed of competing sequence *s*^0^ from scenario *c*^0^, feedforward excitation to co-active assemblies must be reduced.

## 4 Discussion

Using a non-linear rate-based model with discrete and pre-configured assemblies, we provided a proof-of-principle for competition and cooperation between neural activity sequences. The model allowed us to study the dynamics of isolated, competing and cooperating sequences. Characterizing conditions for successful sequence progression, we can attribute specific roles to the interactions within and between assemblies. Projections between subsequent excitatory populations ensure sequence progression. However, if too weak, activity does not propagate, and if too strong, activity saturates. Recurrent excitatory and inhibitory interactions implement balanced amplification which boosts weak excitatory inputs and prevents saturating activity (Murphy and Miller, 2009; Hennequin et al., 2012). Thus, with increasing recurrency, a larger range of excitatory inputs is permissible. Further, the boost of weak inputs is especially beneficial in the competition scenario and allows the surviving sequence to quickly recover. Excitatory interactions between co-active assemblies allow weak sequences to win against a stronger competitor, but such interactions slow the propagation of activity. Shifting feedforward excitation from co-active to subsequent assemblies of cooperating sequences increases sequence speed, enabling successful replay without slowing sequence propagation.

In the case of a single sequence, the analytically predicted minimal feed-forward and recurrent weights are in close agreement to the non-linear rate model. We contrasted the simulation results with an even simpler model of assembly sequence progression, comprised of a linear dynamical system and with only projections to subsequent assemblies (Chenkov et al., 2017). Assuming that the activation time of a preceding excitatory population is much longer than the rise time of the subsequent excitatory population, we derived conditions for sustained activity propagation. For moderate and large recurrent connection strengths, the simulation quantitatively agreed with the analytic prediction. As part of the analytic approximation, we used the same scaling factor as Chenkov et al. (2017). This factor has been fitted to match the lower bounds for sequence progression in a spiking neural network. Thus, the estimated conditions for successful progression should translate to similar dynamics in a spiking neural network.

For weak recurrent interactions, the results of the non-linear network deviate from both the analytic approximation and a previously published spiking neural network (Fig. 1c, Chenkov et al., 2017). Even without recurrent interactions, the non-linear rate model allows single sequence progression for a very narrow range of feed-forward projections. However, in contrast to the analytical approximation and the spiking model, weaker feed-forward interactions are required in the non-linear rate model. A definite explanation requires further investigation. Potentially, the divergence is a result of the different wiring of feed-forward inhibition. In the non-linear rate model, all assemblies, even when in the same sequence, send and receive feed-forward inhibition. In the analytic approximation there is no feed-forward inhibition within a sequence, only between sequences. In the published spiking model feed-forward inhibition is not assembly specific and connection strengths are plastic, making it difficult to compare (Chenkov et al., 2017).

Our results highlight a key constraint on which synapses may be potentiated to support successful pairing of activity sequences. We report that direct excitatory interactions between co-active assemblies lead to increased activation times (Fig. 4) and slower sequence propagation (Fig. 5). Maintaining propagation speed for paired sequences was made possible by potentiating excitatory projections to subsequently active excitatory populations in the cooperating sequence. Such temporally skewed potentiation may naturally occur via asymmetric spike-timing-dependent plasticity during encoding (Klos et al., 2018; Miner and Tetzlaff, 2020). We note that several other mechanisms may modulate speed: Dynamic firing rate adaptation to mimic refractory periods (Wilson and Cowan, 1972), inhibitory oscillations to rhythmically gate propagation (Recanatesi et al., 2015), or inhibitory plasticity to maintain EI balance (Vogels et al., 2011).

The presented results equally relate to the creation of new synapses as well as to potentiation of existing synapses. The strength of an individual connection is defined by the product of population size, the average connection probability and the synaptic weight. Unlike population size which also affects other projections of this population, the specific connection probability and synaptic weight are interchangeable scaling factors.

Weak sequences may also compensate for small assembly sizes by potentiating recurrent interactions, weakening feed-forward inhibition, or recruiting more neurons (assembly outgrowth, see Tetzlaff et al., 2015; Lehr et al., 2022). Here, the underlying learning scenario is highly simplified. We assume that during learning pre-configured, recurrently interacting assemblies are activated by external input. This is thought to induce the formation or potentiation of excitatory projections between subsequently activated excitatory populations. For this reason we only evaluated the possibility that weak sequences compensate for small assemblies by strengthening projections between subsequent excitatory populations.

Competition between neural activity sequences may be directly observed in hippocampal recordings. If reactivation of neural activity sequences in the hippocampus is indeed the outcome of a competition process, signatures of this process should be detectable. In the presented model, competition dynamics are characterized by an initial rise in the activity of assemblies of different sequences, followed by reduced activity due to mutual inhibition, until one sequence starts to out-compete the others. Such dynamics should be particularly strong if competing sequences are of equal strength. Studying the reactivation of place cell sequences after running on two or more distinct linear tracks may be an adequate experimental paradigm (Silva et al., 2015; He et al., 2020).

In summary, our work investigated the interaction of multiple sequences of different strengths within a recurrently connected network. We considered scenarios of competition and cooperation between interacting sequences and characterized the effects on sequence reactivation and sequence dynamics. We showed that pairing weak sequences allows them to win over a stronger competitor. This has implications for hippocampal replay – the number of hippocampal neurons recruited to represent certain types of information strongly differ between sensory modalities (Salz et al., 2016; Danjo et al., 2018), thus making it important to develop a theoretical understanding of how heterogeneity in assembly size influences replay statistics.

### 5 Methods

Simulations and analysis were performed with Jupyter notebooks 6.0.3 and Python 3.7.8 with standard libraries, such as NumPy 1.18.5, SciPy 1.18.5, Matplotlib 3.2.2 and SymPy 1.5.1. All code is available at https://github.com/tristanstoeber/sequence_competition_cooperation.

### 5.1 Assembly sequences in a non-linear rate model

In the non-linear rate model each assembly is formed by one excitatory and one inhibitory population. The evolution of rate, 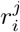, of a given population *i* of sequence *s*^*j*^ is described by

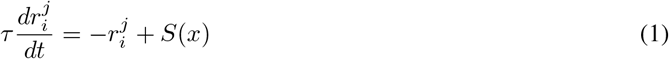

with *τ* a fixed population time constant, equal across all populations.

The sigmoidal activation function *S* over the input *x* is defined by

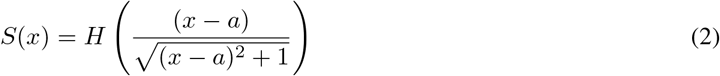

with the Heaviside function *H* (compare Fig. 1a) and *a* = 1 × 10^−7^ a small constant rightward shift of the activation function, preventing numerical imprecision around *x* = 0 from inadvertently driving network activity.

Each population in sequence *s*^*j*^ receives input by recurrent excitatory and inhibitory projections with strengths 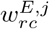 and 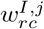 Excitatory populations may receive additional excitatory input by the preceding assembly of the same sequence with strength 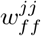 . In the case of cooperating sequences, each excitatory population receives excitatory input of a co-active assembly of another sequence *s*^*m*^ with strength 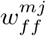. All assemblies send feedforward inhibition to each other, e.g. they excite each others inhibitory population. Thus, in addition to the recurrent input 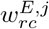 from their associated excitatory population, they receive input 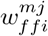 from all remaining *n* excitatory populations 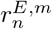 of all sequences *s*^*m*^. Thus, the full input to an excitatory population 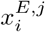 and an inhibitory population 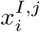 of assembly *i* in sequence *s*^*j*^ is described by:

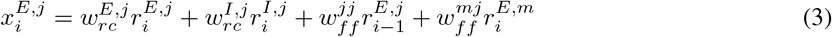

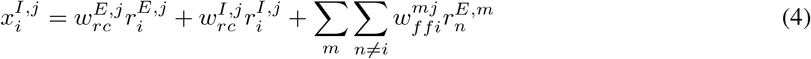

Weight values are the product of the number of excitatory, *M*^*E,j*^, and inhibitory, *M*^*I,j*^, neurons in the sending population, equal for all assemblies in a given sequence *s*^*j*^, as well as recurrent, *p*_*rc*_, feed-forward, *p*_*ff*_, and feed-forward inhibitory, *p*_*ffi*_, connection probabilities and excitatory, *g*^*E*^, or inhibitory, *g*^*I*^, synaptic strengths.

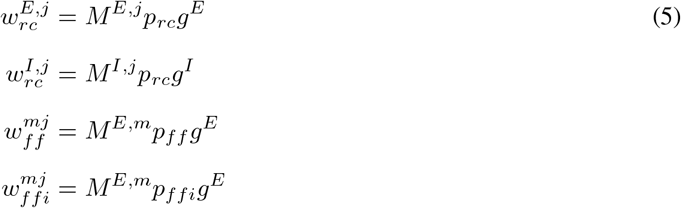

Sequences are comprised of *n*_*ass*_ assemblies, all assemblies within a given sequence are equal in size and the *E/I* size ratio is fixed to *M*^*E*^*/M*^*I*^ = 4.

### 5.2 Simulation and data analysis

Simulations were run for a fixed time interval and a fixed step size with the solve_ivp function in SciPy’s integrate package with integration method LSODA. As initial condition, the excitatory population in the first assembly of each activated sequence *s*^*j*^ is set to 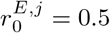, while all other rates are at zero.

To be classified as successfully progressing, a sequence must satisfy the following four conditions: 1) All active: All assemblies must be activated. There should exist at least one point in time during which the activity of a given excitatory population exceeds a minimal threshold *r*_*min*_. 2) All informative: Each excitatory population must exceed the activity of others at least one point in time. 2) Sparse activity: While the sequence is running, maximum firing rates at any given point in time must not be reached by more than two assemblies. To exclude numerical edge cases we consider assemblies to have similar firing rates, whenever the absolute value of the difference is less than *r*_*tol*_. Allowing two assemblies to both have peak activity is necessary for the time points when decreasing activity of the previous and increasing activity of the subsequent assembly are equal. 4) Order: Activation times must maintain sequence order. The order of peak activities agrees with the predefined order of assemblies in the sequence. Given our predefined one-step feed-forward interactions this is almost always the case, though we mention it for completeness.

Sequence speed is determined as the inverse of the median interpeak interval of excitatory populations. Before determine timepoints of peak activation, we rounded values to a precision of *r*_*t*_*ol* and ignored values below *r*_*m*_*in* to avoid numerical fluctuations to be considered a.

For a summarized description of all parameters used in the non-linear rate model see Table 1. For a summary of all used parameters see Table 2.

**Table 1:**
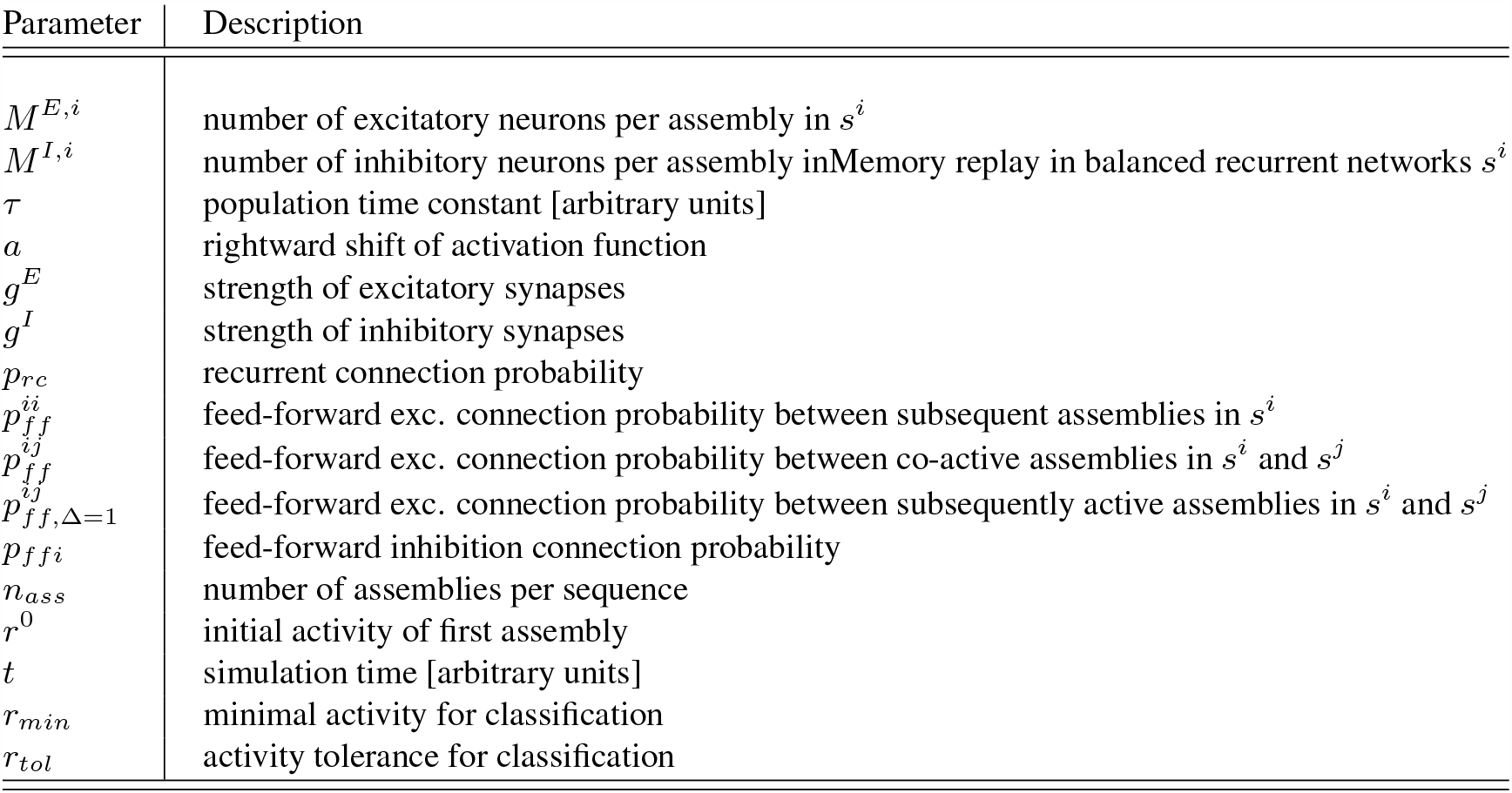
Description of parameters in the non-linear rate model.

**Table 2:**
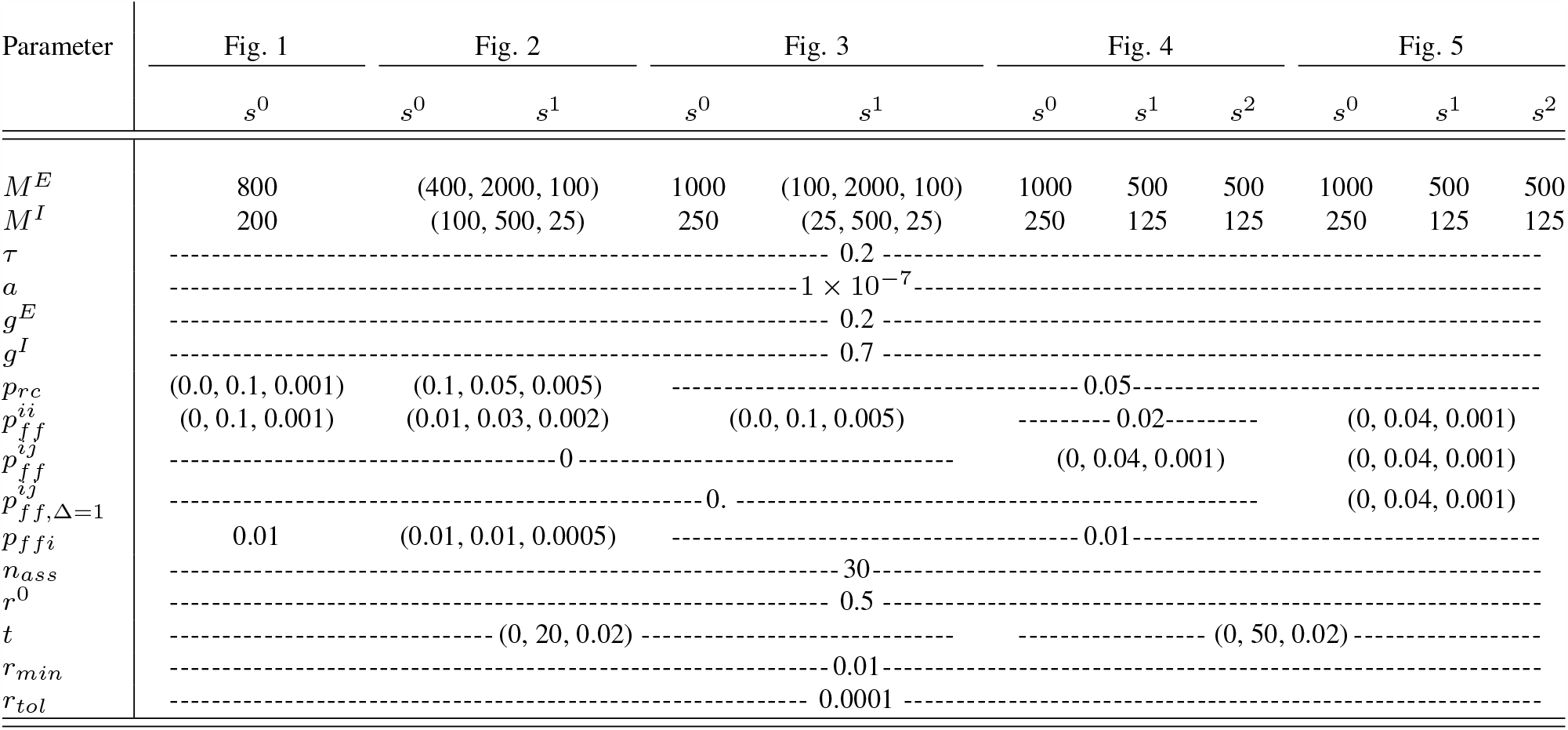
**Parameters of the non-linear rate model** For each figure and sequence. Ranges are indicated with (start, stop, stepsize). Dashed lines indicate values applying to multiple columns.

### 5.3 Linearized approximation of assembly sequence progression

We approximate assembly sequence progression in a linear dynamical system, as in Chenkov et al. (2017). To study both sequence competition and cooperation, we define three sequences: *s*^0^, *s*^1^, *s*^2^. Each assembly in position *i* of sequence *s*^*j*^ is described by the rate of its excitatory 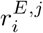 and inhibitory 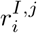 population . We combine population rates in a single vector 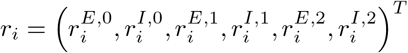 and write the full system as

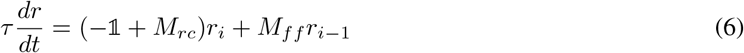

with the unity matrix −𝟙 representing self-dampening, *M*_*rc*_ recurrent interactions and *M*_*ff*_ feed-forward projections from preceding assemblies to the same or other sequences. In each assembly excitatory and inhibitory populations are recurrently interacting (see Fig. 1b). Excitatory recurrent projections between assemblies of sequence *s*^*j*^ are summarized by 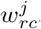, representing the number of participating neurons, connection probabilities and connection weights. Recurrent inhibitory projections, 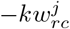, are scaled by factor *k*, the relative strength of inhibition, summarizing both differences in inhibitory populations sizes and synaptic weights. Thus all recurrent interactions are represented by:

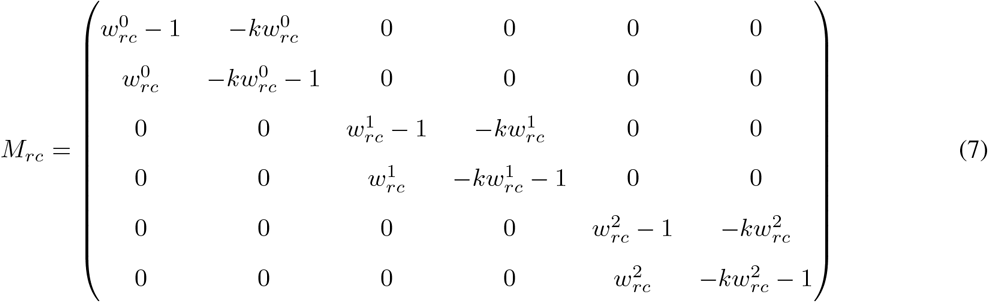

To simplify the mathematical treatment, we model interactions of assemblies within and between sequences only via excitatory feed-forward projections to subsequently active assemblies (Fig. 1b). As such, feed forward projections from sequence *s*^*j*^ to *s*^*m*^ originate from excitatory populations and target either the excitatory or the inhibitory population with strength 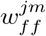 and 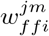, respectively.

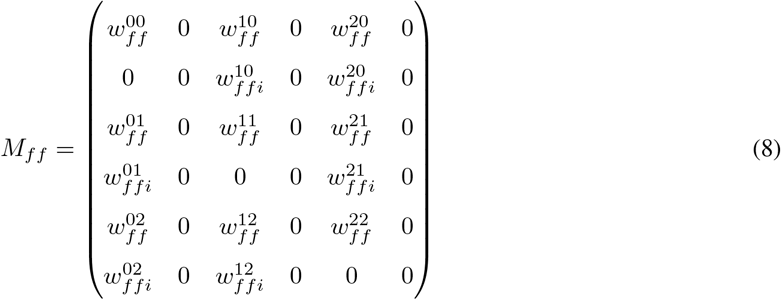

Under the assumption that the activity in the previous assembly persists much longer than the population time constant *τ*, we can consider the steady state 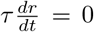 as a sufficient approximation. With this, we can further simplify the system to:

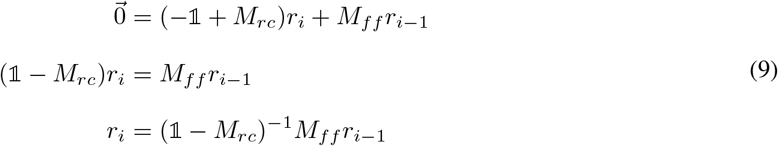

Because inhibitory populations are assumed to have only recurrent projections, we can insert the expression for each inhibitory population 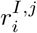 into its respective 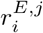 and reduce the system of equations to:

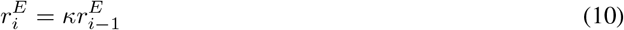

With 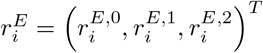 and

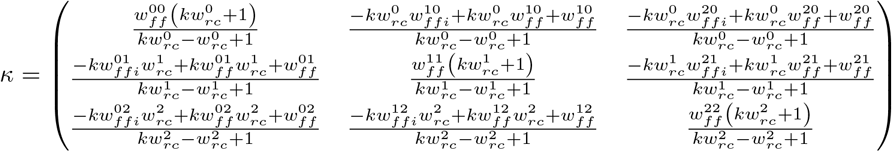

To connect parameters of the population based model to neurons, connection probabilities and synaptic strengths, we proceed as in the non-linear rate model and set recurrent and feed-forward weights to

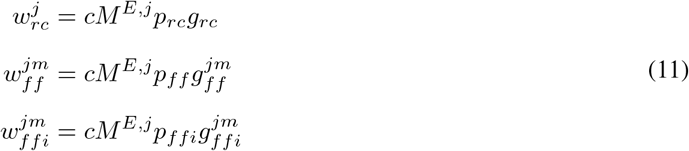

with *c* a scaling parameter related to the slope of the neurons’ input-output transfer function (Chenkov et al., 2017, see below), *M*^*E,j*^ the number of neurons per excitatory assembly of sequence *s*^*j*^, *p*_*rc*_ and *g*_*rc*_, *p*_*ff*_ and 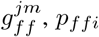 and 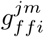, the connection probabilities and synaptic weights for recurrent, feed-forward excitation and feed-forward inhibition, respectively. Further, we assume that the network operates in an approximately balanced state and set *k* = 1.

For the single sequence scenario, we can express the firing rate of 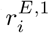 as a function of 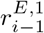

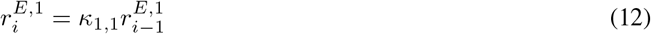

Thus we can express the condition for marginally stable propagation of sequence *s*^1^ as

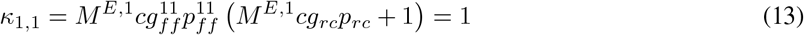

We determine the minimal required recurrent connection probability by solving for *p*_*rc*_.

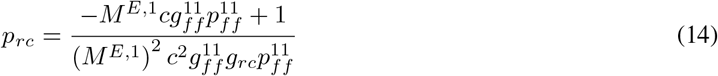

Further, we also derive *c* from equation 13.

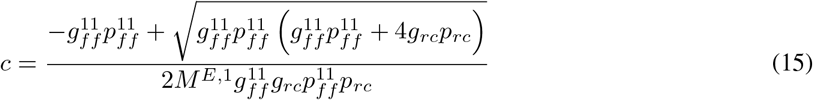

We determine *c* = 0.163 with the parameters of the successful example sequence *s*^2^ from Fig. 1c (*p*_*rc*_ = 0.035, *p*_*ff*_ = 0.01) rounded to the third decimal. We keep *c* at this constant value throughout the article.

To study the relation between required feed-forward weight and excitatory population size in a competition scenario, we add the influence of a competing sequence *s*^0^

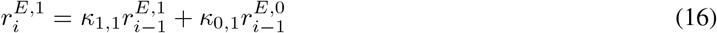

Again we define the condition for marginal stability as

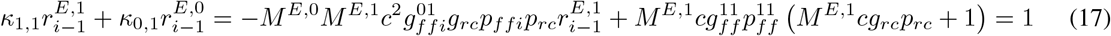

and solve for 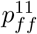

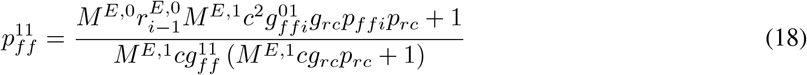

By setting 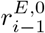 to 1 or 0, we can define a scenario with and without competition.

